# Development of the myzozoan aquatic parasite *Perkinsus marinus* as a versatile experimental genetic model organism

**DOI:** 10.1101/2021.06.11.447996

**Authors:** Elin Einarsson, Imen Lassadi, Jana Zielinski, Qingtian Guan, Tobias Wyler, Arnab Pain, Sebastian G. Gornik, Ross F. Waller

## Abstract

The phylum Perkinsozoa is an aquatic parasite lineage that has devastating effects on commercial and natural mollusc populations, and also comprises parasites of algae, fish and amphibians. They are related to, and share much of their biology with, dinoflagellates and apicomplexans and thus offer excellent genetic models for both parasitological and evolutionary studies. Genetic transformation has been previously achieved for select *Perkinsus* spp. but with few tools for transgene expression and only limited selection efficacy. We thus sought to expand the power of experimental genetic tools for *Perkinsus marinus* — the principal perkinsozoan model to date. We constructed a modular plasmid assembly system that enables expression of multiple genes simultaneously. We developed an efficient selection system for three drugs, puromycin, bleomycin and blasticidin, that achieves transformed cell populations in as little as three weeks. We developed and quantified eleven new promoters of variable expression strength. Furthermore, we identified that genomic integration of transgenes is predominantly via non-homologous recombination and often involves transgene fragmentation including deletion of some introduced elements. To counter these dynamic processes, we show that bi-cistronic transcripts using the viral 2A peptides can couple selection systems to the maintenance of the expression of a transgene of interest. Collectively, these new tools and insights provide new capacity to efficiently genetically modify and study *Perkinsus* as an aquatic parasite and evolutionary model.

## INTRODUCTION

*Perkinsus* species are major marine parasites that cause disease and mortalities in commercially and environmentally important marine molluscs including oysters, abalone, clams and other shellfish (Choi and Park, 2010; Villalba et al., 2004). *Perkinsus marinus* notably causes disease in *Crassostrea virginica* (Eastern Oyster) and has caused mass mortalities and major impact on fisheries since the 1950s (Smolowitz, 2013). The parasite is taken up by the filter-feeding host animal and phagocytosed by host hemocytes mediated by a galectin receptor (Lau et al., 2018; Tasumi and Vasta, 2007). Once inside, the parasite proliferates in a parasitophorous vacuole as it acquires nutrients and evades host defensive oxidative stress (Schott et al., 2019). Upon replication and then egress from the host cell, the released parasites continue to infect more hemocytes or proliferate directly within the host haemolymph. Ultimately the host tissues are overwhelmed, and parasites are released by the moribund animal into the water column where they can infect further shellfish. There are currently no practical treatments available to limit the spread of the parasite and *Perkinsus* infection is listed on the World Organisation of Animal Health (OIE) as a reportable disease of molluscs. In addition to the impact on commercial fisheries, molluscs are major environmental engineers owing to their prodigious water filtration and positive effect on water quality (Smolowitz, 2013). The wide host range of *Perkinsus* spp., and their increasing prevalence in response to global ocean temperature changes, make them critically important parasite taxa (Cohen et al., 2018). Moreover, other members of the Phylum Perkinsozoa are parasites of microeukaryotes and are responsible for controlling major algal blooms (Alacid et al., 2017; Norén et al., 1999), and in recent years others have been recognised as parasites of other animals including fish and amphibians (Chambouvet et al., 2015; Freeman et al., 2017).

The perkinsozoa also provides a very relevant lineage for understanding the evolution of other major eukaryotic groups, namely apicomplexans and dinoflagellates, with which they form the clade Myzozoa. Phylogenetically, Perkinsozoa is the most basal lineage to dinoflagellates after their divergence from the sister apicomplexan group (Bachvaroff et al., 2013; Janouškovec et al., 2017; Saldarriaga et al., 2003). Prior to molecular phylogenies, however, *Perkinsus* was categorised as a member of the Apicomplexa as morphological studies revealed that they possess an apical complex, the characteristic invasion structure of this ubiquitous animal parasite group (Perkins, 1976). Their subsequent phylogentetic assignment to the Dinozoa is consistent with *Perkinsus* sharing some, but not all, of the genomic peculiarities of dinoflagellates (Perkins, 1996; Zhang et al., 2011). This mixture of traits is indicative of Perkinsozoa’s intermediate phylogenetic position. Further, *Perkinsus* has retained a plastid but lost photosynthesis independently of both dinoflagellates and apicomplexans and, furthermore, they have independently developed parasitic lifestyles (Robledo et al., 2011). Thus, *Perkinsus* shares common biology of both groups including a propensity for convergent adaptations. While genetic tools are available for some apicomplexans, these tools are very limited or unavailable in dinoflagellates (Chen et al., 2019; Nimmo et al., 2019). This further positions *Perkinsus* as a key model for understanding the biology of Myzozoa.

In 2008 *Perkinsus marinus* was genetically transformed for the first time by expression of a GFP reporter protein fused at the C-terminus of a highly expressed predicted cell surface protein called MOE under control of its endogenous regulatory sequences (Fernández-Robledo et al., 2008). This achievement heralded an exciting new era for experimental biology in *P. marinus*. Since then, drug selection systems have been demonstrated to be feasible, albeit with selection times of multiple months (Sakamoto et al., 2016, 2019). We have sought to improve the accessibility of methods for *P. marinus* transformation and extend the efficiencies and capabilities of these methods to better understand myzozoan biology in this economically and evolutionarily important lineage.

## RESULTS

In pursuit of a simple transformation method without the use of proprietary reagents, we adopted a defined transformation solution, ‘3R’, used also for *Trypanosoma* spp. electroporation (Faktorová et al., 2020). This substitution provides both inexpensive and efficient (see below) transformation. We have also tested transformation methods other than electroporation. While we found chemical-based transformation reagents were ineffective or not tolerated (not shown), we previously showed that glass abrasion methods resulted in transformation efficiency of up to 1%, alleviating even the need for an electroporation device to transform *P. marinus* (Faktorová et al., 2020).

### Simultaneous expression of multiple transgenes

To date only single coding sequences have been expressed in *P. marinus*, typically as fusion proteins (Cold et al., 2017; Fernández-Robledo et al., 2008; Sakamoto et al., 2016, 2019). To allow for further development of genetic modification methods we asked if multiple genes could be expressed simultaneously from a single expression vector. To do this we exploited the Golden Gate modular plasmid assembly method modelled on the ‘MoClo Plant Took Kit’ (Fig. 1A) (Engler et al., 2014; Werner et al., 2012). ‘Level 0’ plasmids were prepared containing basic genetic modules (promoters, terminators, coding sequences) flanked by Type IIS restriction enzyme sites with adjacent specific overhangs to direct the order of module assembly. These Level 0 modules were assembled as ‘scarless’ (lacking the original restriction enzyme recognition site) expression units into ‘Level 1’ acceptor plasmids, again with adjacent Type IIS restriction enzyme sites. The Level 1 plasmids are specific to the gene position order of the final ‘Level 2’ assembly product where up to seven expression units can be simultaneously assembled or capped with an ‘end linker’ unit if less than seven are required. Using this method, we generated several plasmids with two, and one with three, expression units. To test of this strategy, we sought to express separately the reporters mCherry and luciferase. When transformed in *P. marinus* cells, both mCherry fluorescence and luciferase-mediated illumination were detected (Fig. 1B). In this case identical promoter and terminator regions were used for both genes, demonstrating that this is not an impediment to multi-gene expression. All multigene expression plasmids proved successful (see below), and the Golden Gate assembly pipeline provided an effective method for their assembly.

**Figure 1.**
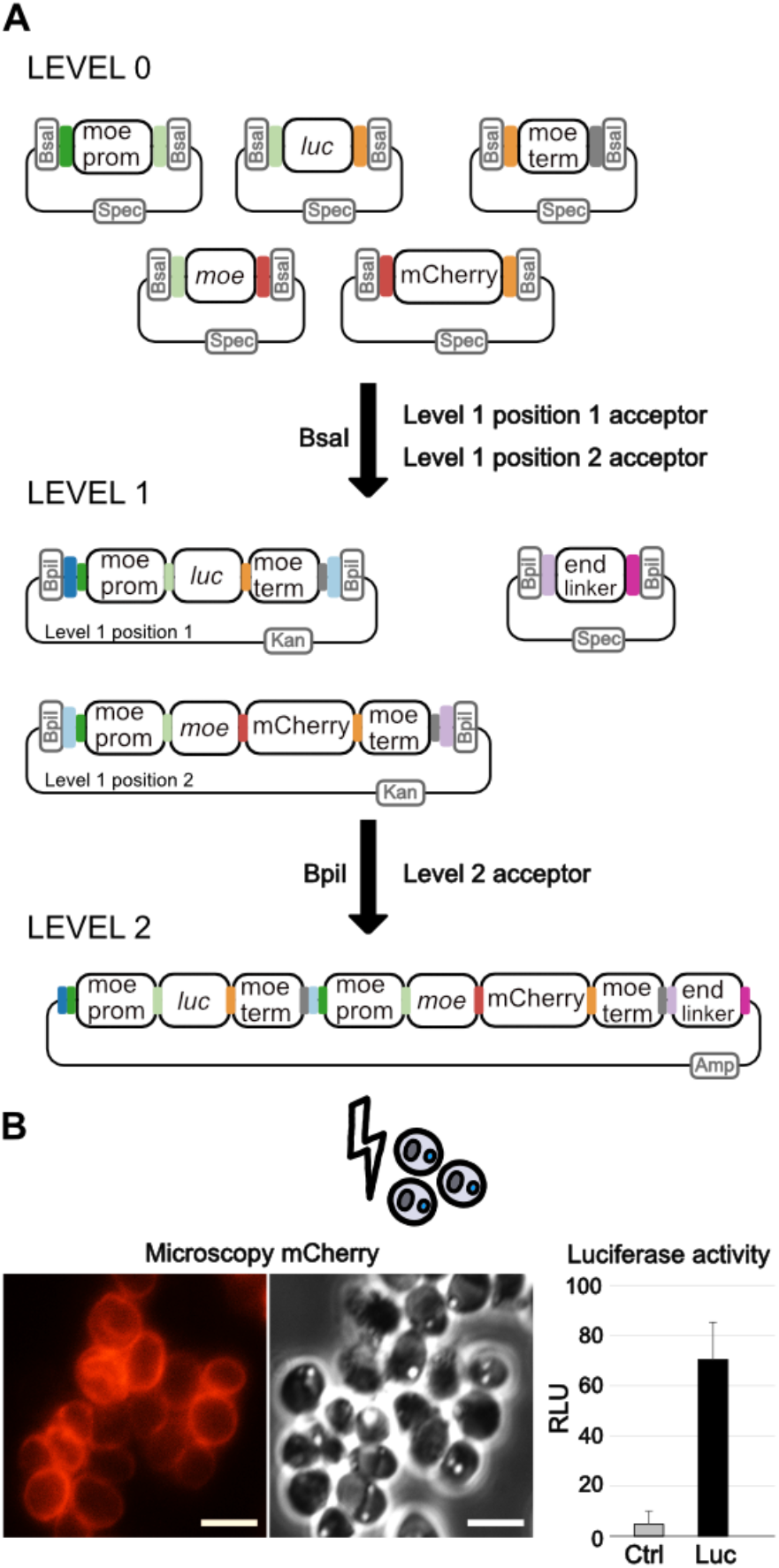
Golden Gate assembly of *P. marinus* transformation plasmids to express multiple genetic elements. (**A**) Level 0 plasmids contain elements such as promoters, coding sequences, and terminators. In this example, we show the previously described *moe* promoter and terminator, the *moe* gene, luciferase, and mCherry as a C-terminal fusion coding sequence. *BsaI* restriction sites generate unique 4-nucleotide overhangs (shown by different colours) that coordinate ordered assembly into a range of Level 1 plasmids. Level 1 expression modules are, in turn, assembled in order into Level 2 acceptor plasmids by coordinated *BpiI* unique overhangs. (**B**) *P. marinus* cells transfected with the Level 2 plasmid show both Moe-mCherry peripheral fluorescence and luciferase activity. Non-transformed cells served as control in the luciferase assay. RLU, Relative Light Units. Scale bars = 5 μm. Error bars = standard deviation.

### Improved selection of transformants

The first transformations performed on *Perkinsus* lacked selection methods and relied on the relatively high efficiency of plasmid uptake to determine the outcomes of transgene expression (Fernández-Robledo et al., 2008). The subsequent expression of resistance genes for bleomycin and puromycin demonstrated the scope for drug-based selection in *Perkinsus*, however, these studies required several months of selection before healthy drug-resistant cells were acquired (Sakamoto et al., 2016, 2019). We sought methods to improve the speed and efficiency of selection of transformed *Perkinsus* cells.

We first exploited flow cytometry to both quantify transformed cell populations where fluorescence reporters were used and as a fast selection strategy to sort them into populations of transformed cells, or single cells for clonal propagation. Using flow cytometry analysis of GFP or mCherry expression we determined that initial transformation efficiencies typical range from 0.001 to 5% (e.g., Fig. 2A). Fluorescence microscopy of transformed cells before and after Fluorescence Activated Cell Sorting (FACS) showed that FACS provided a highly efficient method of selection (Fig. 2B). *P. marinus* was also amenable to single cell sorting and subsequent generation of clonal cell populations (see examples below).

**Figure 2.**
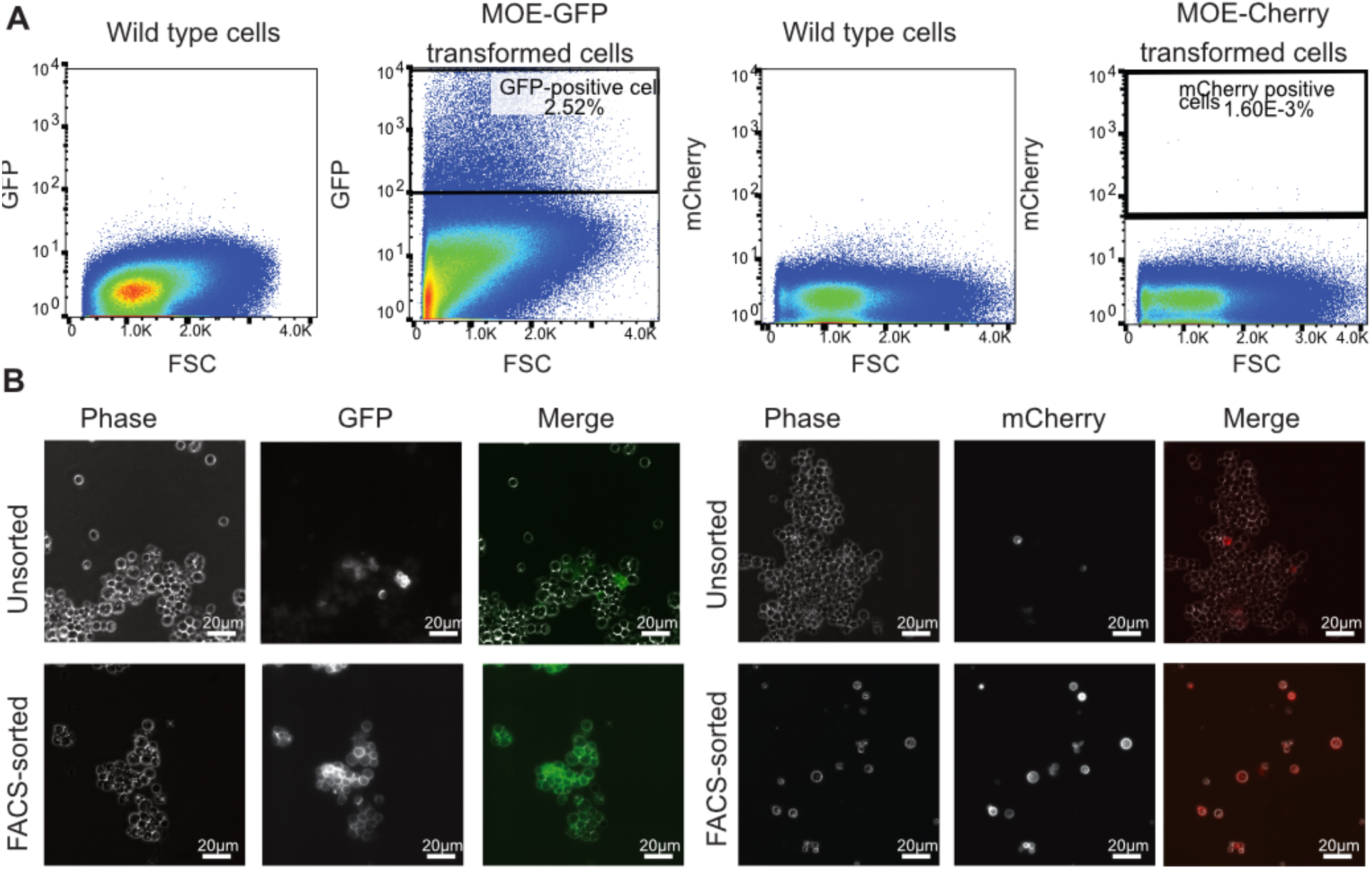
Flow Cytometry to assessment and sort transformed *P. marinus*. (**A**) FACS bivariate plot showing forward scatter (FSC) and fluorescence intensity (eGFP or mCherry). Conservative gates distinguish untransformed and positively fluorescent cells. (**B**) Fluorescence microscopy of cell populations pre- and post-sorting showing effective enrichment of transformed cells. Scale bars 20 μm.

Despite the demonstrated utility of FACS for selection of *Perkinsus* cells transformed with a fluorescence marker, the relatively high cost of FACS facilities and the utility of non-fluorescent reporters means that it is not suitable for all experiments. Drug-selection for transformant cells is a widely used method for most established model organisms. We speculated that the previous application of drug selection in *Perkinsus* was slow and inefficient due to suboptimal culture conditions during selection, with no change of medium or selection drug. To test this, we designed a selection regime that regularly replenished both (Fig. 3A) and tested this using three drugs (bleomycin, puromycin and blasticidin S) and their corresponding drug resistance genes as selectable markers. We first verified the sensitivity of *P. marinus* to each of these drugs over 15 days of cell growth (Fig. 3Bi). Upon transformation with plasmids containing each drug resistance gene and a MOE-mCherry fluorescent reporter fusion, cells were allowed to recover for 3-5 days. Successful transformation of a subpopulation of cells was confirmed using fluorescence microscopy for MOE-Cherry, and then bleomycin (200 mg/ml), puromycin (20 μg/ml), or blasticidin S (200 μg/ml) were added to commence selection. Over the next three weeks, once a week the parasites were gently pelleted by centrifugation and resuspended in fresh medium and drug. At three weeks, healthy growing cells capable of 50% serial passaging were evident and flow cytometry showed strong mCherry fluorescence in this population (Fig. 3C). These positively selected cells were then tested for drug-resistant growth rates (Fig. 3Bii). Cells transformed with bleomycin and puromycin resistance genes showed equivalent growth rates in drug comparted with drug-free medium, including at a two-fold increase in drug concentration to the selection conditions. While the blasticidin S selection regime was also successful, resistant cells grew slightly slower in drug compared to without drug. To test if prolonged selection beyond 3-4 weeks was required to purify the transformed population from untransformed cells, flow cytometry was used to track the distribution of MOE-mCherry fluorescence in the cell populations under drug selection with blasticidin S and puromycin (Fig. 3C). The fluorescence profile at 3-4 weeks of drug selection did not further change at 7-8 and 11-12 weeks. Our data demonstrate that effective selection of transformants can be achieved in only 3-4 weeks with our selection regime, and this works well both for previously used drugs bleomycin and puromycin, and for the additional selection agent blasticidin S. The subsequent experiments described in this report all used this new selection regime.

**Figure 3.**
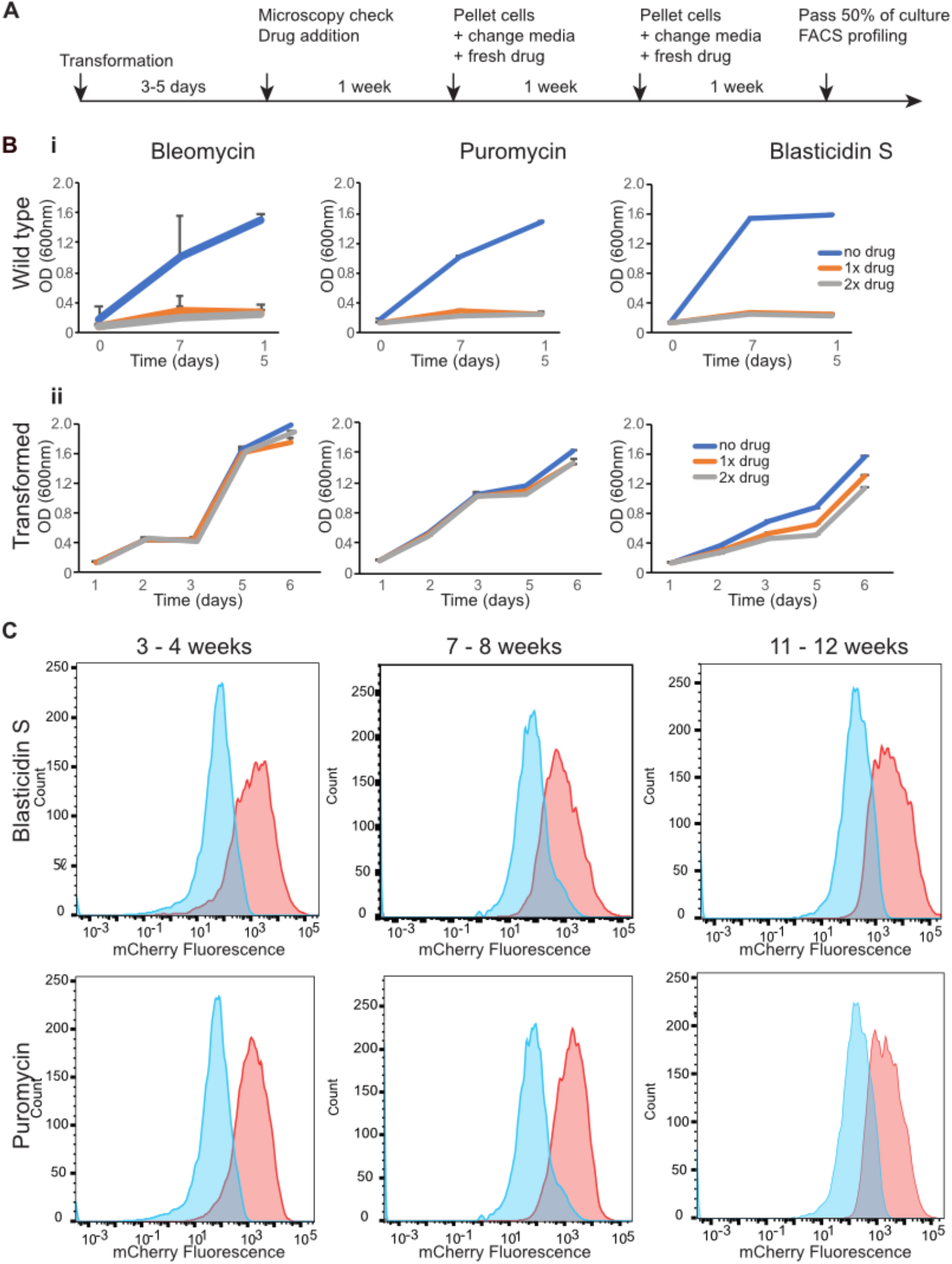
Efficient drug-selection of transformed *P. marinus*. (**A**) Scheme of the drug-selection regime including medium and drug replenishment. (**B**) Drug sensitivity of i) untransformed cells and ii) drug-resistance gene-transformed cells during growth assays in 0x (blue) 1x (orange) or 2x (grey) selection-strength concentrations of drug (bleomycin 200 μg/ml, puromycin 20 μg/ml and blasticidin S 200 μg/ml). Cell growth was measured by optical density at 600nm. (**C**) FACS plots (cell counts versus mCherry fluorescence) of non-drug selected (blue) and drug-selected (orange) populations of transformed cells over three selection time periods.

An alternative route to stable transgene expression is the inclusion of plasmid maintenance elements. The yeast artificial chromosome element CEN6-ARSH4 has been shown to increase the retention of episomal plasmids in diatoms (Diner et al., 2016; Karas et al., 2015). To test if this element could promote plasmid retention in *Perkinsus*, cells were transformed with one or two MOE-mCherry reporter plasmids differing only by the presence or absence of the CEN6-ARSH4 sequence (Fig. S1A). Transformed cells were FACS sorted for mCherry-expressing cells three days post transformation, and then allowed to grow unselected for a further 14 days. Over this time most cells lost fluorescence and there was no significant difference between those with or without CEN6-ARSH4 (Figure S1B).

### Determination of transgene genome integration events

To further develop the applications and interpretation of transformation in *Perkinsus* it is necessary to understand the genomic outcomes of the transformation events. We have sought to characterise these events by whole genome sequencing of transformed *P. marinus* to ask if, and how, transgenes might be integrated into the nuclear genome. From four clonal cell lines derived by single-cell FACS sorting, genomic sequencing identified that transgenes were integrated into the genome in a variety of manners (Fig. 4). No plasmids were integrated in their entirety, rather fragments of the plasmids integrated that corresponded to the mode of cell selection: either fluorescence or bleomycin resistance. In one case only the drug resistance gene (*ble*) was integrated, in another only the fluorescent reporter protein. In another case multiple integrations of a single plasmid were seen. On three separate occasions, integration was seen at sites proximal to retrotransposable elements. We found no cases of integration of our plasmids in at the endogenous MOE locus, despite up to several kilobases of MOE coding and regulatory sequences in some plasmids. This suggests that homologous recombination machinery was not involved in plasmid integration. Together our genomic data verify that stable transformation is the result of genome integration of transgenes. Disruption of the integrated cassette, however, can lead to the loss of elements of these constructs including transgenes that might not be under selection.

**Figure 4.**
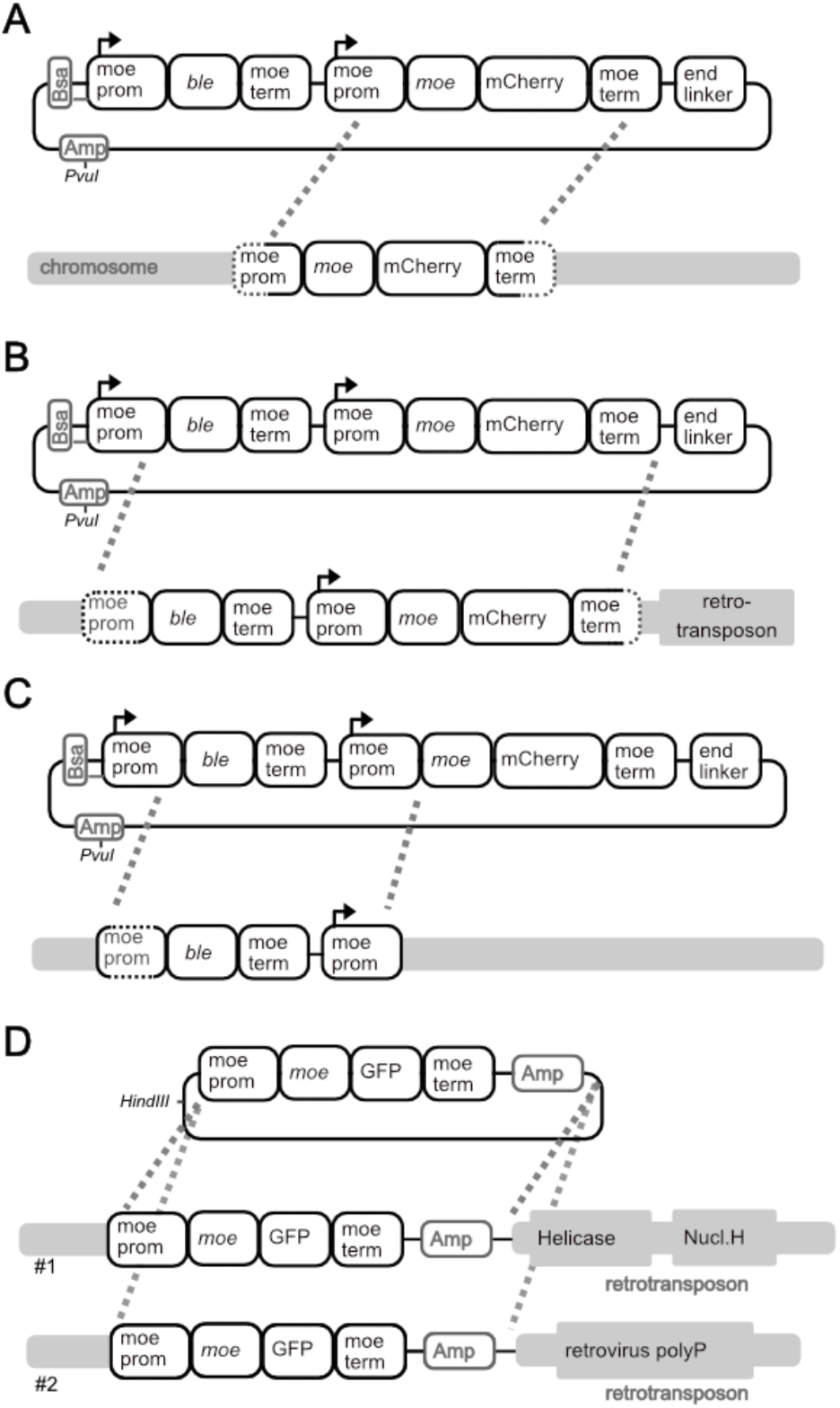
Integration of plasmid genetic elements into *P. marinus* chromosomes (grey) determined by whole-genome sequencing. Five integration events were seen over four clonal cell lines (**A-D**) derived by single-cell FACS sorting. Proximity to retrotransposable elements (grey boxes) of some integration events is indicated. Dashed lines indicate plasmid insertion points. Restriction enzyme sites (*PvuI, HindIII*) indicate where the plasmids were linearised prior to transformation (1:1 circular:linear plasmid mixes were used for transformations). Short-read sequence assembly data of integration sites are available at DOI: 10.6084/m9.figshare.14611578.

### Expansion of functional promoter sequences and transgene expression stability

To date, all reported transformation of *Perkinsus* used the 5’ upstream sequence of the highly expressed transcript for the MOE protein to drive protein expression (Cold et al., 2017; Fernández-Robledo et al., 2008; Sakamoto et al., 2016, 2019, 2021). We tested for further useful promoter sequences that might enable a range of levels of expression of transgenes. To identify candidate promoters, we used a recently generated improved genome assembly for *P. marinus* and RNA-seq data from log-phase cells to provide insight into gene transcriptional expression profiles (Gornik, Flores, Gupta and Waller, unpublished). From these datasets we selected 5’ upstream sequences from ten genes that encode proteins with a known function (Fig. 5A). The candidate promoters were fused upstream of the luciferase coding sequence and after approximately five weeks of puromycin selection luminescence was measured (Fig. 5B, E). The MOE promoter was used as a positive control and benchmark for luciferase expression. Eight of the promoter regions tested showed significant luciferase activity compared to the un-transformed cells (as negative control) with a range of expression strengths compared with the MOE promoter. A further three (H4, SMC, and Sec61) showed low but variable expression. The two shortest promoter sequences tested (GAPDH and HSP90) showed the highest expression. Some variation in expression level was seen between independent replicate transformations, which might reflect difference in the integration events for the expression plasmids. Some of the intergenic sequences that we tested have upstream genes on the reverse strand, so we tested for bidirectional promoter activity for one of these sequences (PRDX). We flanked the ‘PRDX’ promoter region with luciferase on its 5’ side and puromycin N-acetyltransferase on its 3’ sides (Fig. 5C). After drug selection of transformed cells, positive luminescence measurements showed that this sequence drives transgene expression in both directions (Fig. 5E, panel ‘fPRDXp’ ‘rPRDXp’).

**Figure 5.**
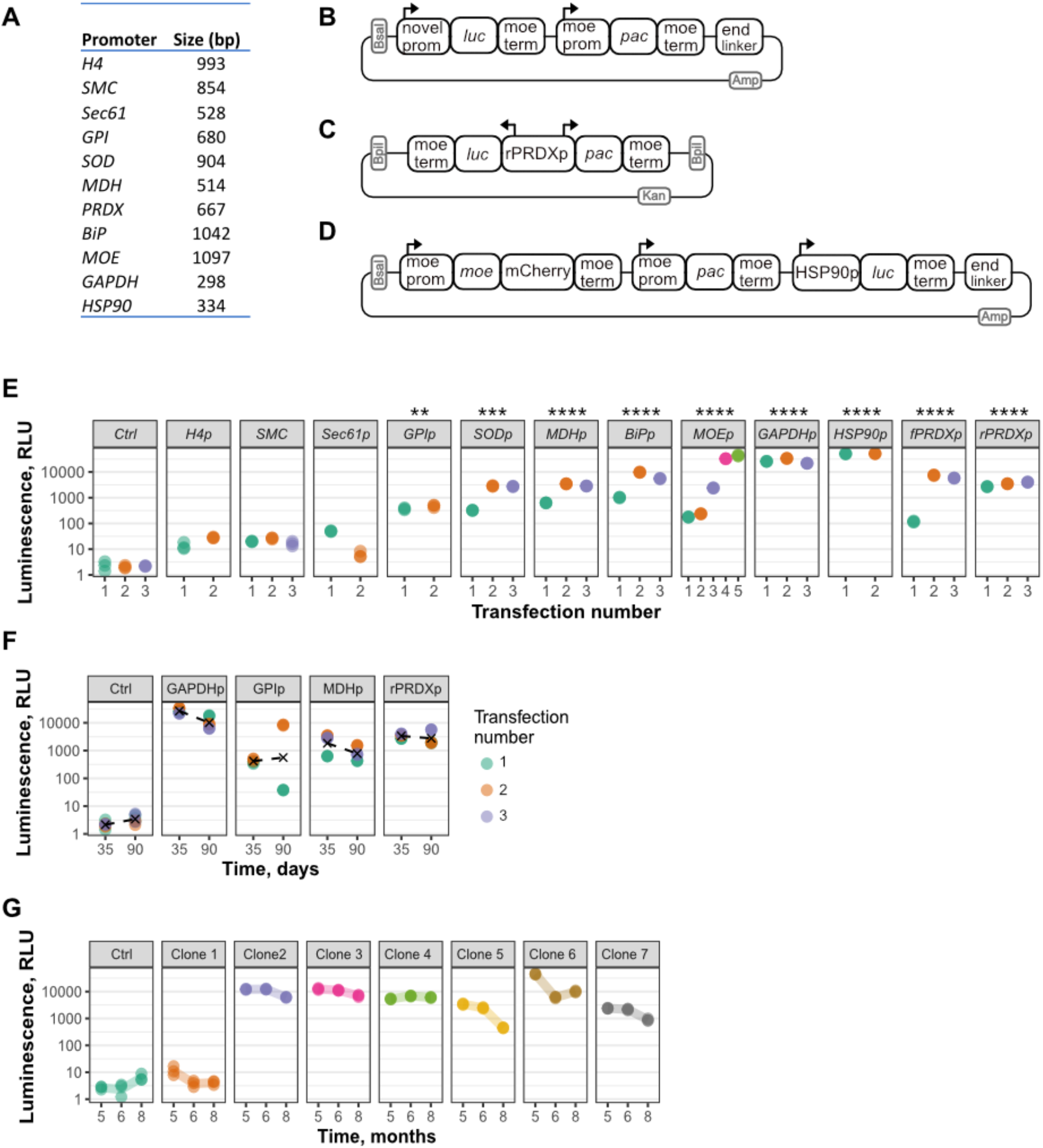
Alternative promoters give different expression strengths. (**A**) Ten new putative promoter sequences, plus the *moe* promoter, of varying size that were assessed for expression strength by luciferase activity. (**B**) Design of expression plasmid where each promoter was separately fused in front of the luciferase coding sequence and combined with a *pac* resistance cassette in a Level 2 plasmid. (**C**) Expression plasmid map with a putative bidirectional promoter region (PRDX) to test for luciferase expression in the reverse direction (rPRDX) compared to that used in B with the *pac* gene expressed in the forward direction from this promoter sequence (fPRDX). (**D**) A three gene expression plasmid with the HSP90 promoter driving luciferase expression. (**E**) Relative Light Unit (RLU) measurements of independent transformations (different colours, with replicate readings plotted to indicate technical variation) for each plasmids of design shown in B and C measured 35 days post transfection. Luminescence values shown as log-transformed medians of each promoter and statistically tested against the non-transformed control cells in a one-way ANOVA analysis of variance, followed by Dunnett’s multiple comparisons test. P-values \*\**P* < 0.01, \*\*\**P* < 0.001, \*\*\*\**P* < 0.0001. (**F**) Stability of luciferase expression in mixed transformation populations measured 35 and then 90 days post transfection. X indicates replicate mean. (**G**) Clonal variation and stability of luciferase expression determined by transformed cells with tri-gene plasmid (**D**) cloned by single cell FACS sorting on mCherry three months post-transformation and luciferase monitored for up to 8 months after transformation. Abbreviations of gene names; Histone H4 (H4), chromosome segregation protein (SMC), Glucose-6-phosphate isomerase 2C cytosolic (GPI), iron-dependent superoxide dismutase (SOD), malate dehydrogenase (MDH), peroxidioxin 2 (PRDX), Binding immunoglobulin protein (BiP), Glyceraldehyde 3-phosphate dehydrogenase (GAPDH) and Heat shock protein 90 (HSP90).

We next asked if transgene expression is stable during prolonged drug selection. For four of the promoter sequences (GAPDH, GPI, MDH, rPrDX), luciferase activity was assayed both at 35 days post-transfection (as in Fig. 5E) and then after an additional 55 days of culture growth luciferase expression throughout despite being puromycin resistant and having been FACS selected for mCherry expression. These data suggest general stability of integrated expression cassettes but are also consistent with cases of partial cassette loss (e.g of the luciferase expression module in clone 1).

### Expression of bi-cistronic expression cassettes using 2A peptides couples drug-selection to expression of transgenes

Given our observations that expression constructs can be disrupted and only partially maintained and/or expressed after integration in *Perkinsus* — observed here both by genomic sequencing and the loss of expression of one out of multiple introduced reporter proteins — we sought a strategy that directly couples the expression of a reporter protein with the selectable marker. The short viral 2A peptides allow expression of multiple proteins from the same regulatory elements (promoter and terminator) and from the same transcript. The 2A sequence mediates a polypeptide break during translation resulting in two protein products (Shaimardanova et al., 2019). We reasoned that linking the expression of a selectable marker downstream of a protein of interest (3’ end of the transcript) would ensure the expression of both proteins when selection is maintained (Fig. 6).

**Figure 6.**
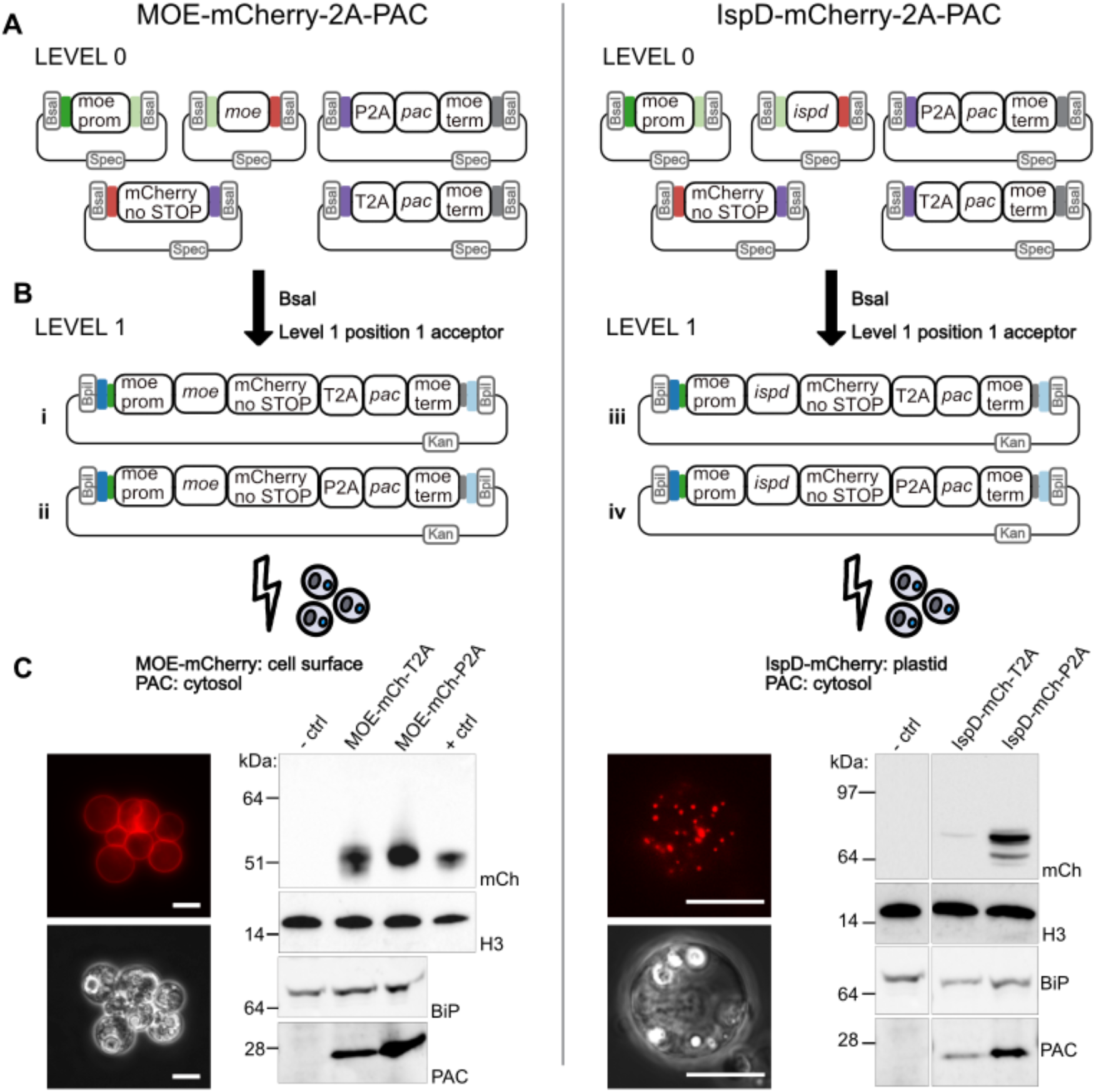
2A peptides allow two proteins to be expressed from one gene. (**A**) A modified Golden Gate assembly design with Level 0 modules allowing the 2A-resistance gene module (*pac*) to be fused directly downstream of a C-terminal reporter protein (mCherry lacking a stop codon). (**B**) Different Level 1 expression plasmids using either a T2A (i, iii) or P2A (ii, iv) skip peptide to co-express a reporter tagged protein of interest (MOE or IspD) and a drug selectable marker (PAC). (**C**) Expression of both proteins observed by fluorescence microscopy and Western blot. Expected size of MOE-mCherry-2A 42.4 kDa, IspD-mCherry-2A 81.7 kDa and PAC 22 kDa. – ctrl, untransformed cells; + ctrl, MOE-mCherry single gene expression. Loading controls: H3, histone H3; BiP, ER chaperone. Scale bar = 10 μm.

To test the utility of 2A peptides in *Perkinsus* we modified our Golden Gate strategy to include two new Level 0 parts that can be fused sequentially downstream of the gene of a interest: 1) the coding sequence of a C-terminal reporter fusion (e.g., mCherry) without a termination codon; 2) the coding sequence of a 2A peptide (either P2A or T2A) linked to the *pac* gene and followed by the *moe* terminator sequence (Fig. 6A). This allowed assembly of a bi-cistronic vector (Fig. 6B) that, when transformed in *P. marinus* and selected on puromycin as described above, resulted in mCherry-positive cells. We tested this strategy for both the surface protein MOE and a predicted plastid protein involved in isoprenoid precursor biosynthesis, 4-diphosphocytidil-2c-methyl-D-erythritol cytidylyltransferase (IspD) (Matsuzaki et al., 2008). Red fluorescence was seen according to the expected cell location of each: the cell periphery for MOE, and punctate structures for the membrane-bound plastid organelles (Fig. 6C). To verify that the 2A-linked transcripts are indeed translated as separate (Fig. 5F). In general, for all cases, ongoing luciferase activity was seen but with variation in independent replicates both up and down. To test if this variation was either due to changing population structure in these mixed transformation cultures, or change with time of individual cell lineages, clonal transformed cell lines were generated. We used a plasmid that separately expressed three proteins: MOE mCherry for FACS sorting, PAC for drug selection, and luciferase under the control of the HSP90 promoter for expression quantitation (Fig. 5D). Clonal cell lines were generated by single-cell FACS sorting based on mCherry expression and thereafter cultured with drug-selection until sufficient cell numbers allowed for luminescence measurements (Fig. 5G). Six of seven clones selected showed strong luciferase expression over three months of testing. A seventh clone showed negligible proteins we analysed the drug resistant cell populations by Western immunoblotting. We detected both MOE-mCherry and IspD-mCherry fusion protein products of expected sizes lacking the PAC protein and, separately, the PAC resistance protein. No detectable higher molecular bands were observed that correspond in size to a single fused protein, indicating that the 2A sequences are efficient at breaking the polypeptide chain during translation in this system. Difference in expression strength is seen for the two IspD fusions and this might indicate integration differences as were seen for other transformants. A weak protein signal of a slightly larger IspD-mCh-P2A species is seen, and this is consistent with the observation of plastid targeted proteins in related apicomplexan taxa where removal of the N-terminal plastid targeting peptides is slow (Waller et al., 1998, 2000). Minor small bands for the IspD cells indicate some level of proteolysis of this fusion. These data demonstrate that 2A peptides are functional in *Perkinsus* and provides an alternative method of expressing multiple proteins from a single vector.

## DISCUSSION

In pursuit of further developing *Perkinsus marinus* as an experimental genetic model system relevant to myzozoan biology we have developed cheap alternative methods to introduce transgenes, and a versatile modular DNA assembly platform for generating transformation vectors that allow simultaneous expression of multiple different genetic elements. We have optimised drug selection regimes that enable rapid generation of transformants and shown that bi-cistronic expression units can couple selection to reporter expression. Moreover, we have expanded the repertoire of available promoters from one to ten and these offer a wide range of expression levels and even one that is bidirectional. We have also determined some of the outcomes of transgene genomic integration in *P. marinus* and these both inform our interpretation of transformation results and design of experiments.

Generally, there are two possible outcomes of transformation with plasmids: 1) maintenance of the plasmid as an episome or, 2) integration of all or part of the plasmid into the genome. Genomic integration can be driven by either homologous or non-homologous DNA repair machinery. Our genome sequencing data show that genomic integration does occurs. While we cannot eliminate the possibility of some plasmids being maintained, the observed loss of one transgene phenotype but not another from transformants (e.g. luciferase but not drug resistance) suggests that the introduced free plasmids are ultimately lost. Moreover, the use of the CEN6-ARSH4 unit that confers episomal plasmid maintenance in other systems did not show evidence of enhanced plasmid maintenance in *P. marinus*. This might indicate divergence of genetic inheritance systems in this taxon that are otherwise conserved in taxa as distant as fungi and diatoms (Diner et al., 2016; Karas et al., 2015). Where we have characterised genomic integration, it is more consistent with non-homologous end joining repair after partial degradation of the plasmid DNA. The lack of evidence of homologous integration at the *moe* locus (albeit with only five integration events characterised) suggests that this repair machinery is not favoured in this system. It is, however, conceivable that any integration at the *moe* locus of our transgenes could have resulted in a negative phenotype, although this seems unlikely given that co-expression of the reporter tagged versions of MOE is tolerated. Furthermore, recently the delivery of Cas9-sgRNA complexes to target the *moe* locus do result in homologous integration of donor DNA molecules (Yadavalli et al., 2021). Thus, it seems that both integration routes are possible, and Cas9-mediated DNA nicking might be required to favour the homologous recombination route. Our observation of three of five integrations occurring in close proximity to identifiable retrotransposon sites indicates that these parts of the genome might be especially available or amenable to non-homologous integration events. Alternatively, transfected episomal plasmids may also integrate in the wake of a retrotransposition of an active retrotransposon in a similar way to that observed and engineered in plants (Vives et al., 2016).

Heterogeneity of the level of transgene expression in cells was often observed, both by microscopy and FACS expression profiles. This might indicate some cell cycle differences in expression during *P. marinus* rapid but non-synchronous growth. It is also possible that differences of genomic integration sites might result in different expression levels. However, we also see some differences in expression in clonal lineages over time. Thus, it might be that ongoing transgene rearrangements occur within the genome, or possibly that epigenetic changes associated with the integration locus occur. The observed occasional loss of expression of one transgene overtime (e.g. luciferase) when only the second is selected for (e.g. drug resistance) provides further evidence of ongoing genomic dynamics. With this in mind, the use of 2A peptides provides a robust way to apply ongoing selection of the expression of a protein of interest, and this system has been recently independently corroborated in *Perkinsus* (Sakamoto et al., 2021). Moreover, we show that these bi-cistronic transcripts can direct their separate proteins to two different locations in the cell: e.g. the resistance PAC protein to the cytoplasm and the reporter fusion protein to the small non-photosynthetic plastid of these organisms. This creates a versatile system, particularly when the expression of the reporter protein might be otherwise difficult to detect.

Recently multiple laboratories have demonstrated an interest in, and the applicability of, *Perkisus* spp. as experimental models (Faktorová et al., 2020). Moreover, the amenability of *Perkinsus* to Cas9-induced homologous recombination has now been demonstrated as well as *Perkinsus* colony growth on solid agar media (Cold et al., 2016; Yadavalli et al., 2021). With the tools that we have generated here, *Perkinsus* is now primed as a powerful and readily accessible genetic model. This offers tremendous scope to test hypotheses of its interaction with and pathogenesis of marine molluscs, and to explore the evolutionary trajectories in cell biology seen more widely in myzozoan protists.

## ACKNOWLEDGEMENTS

We are grateful to Mark Carrington for providing the BiP antibody, Eelco Tromer for technical assistance with codon-optimisation of construct parts, Konstantin Barylyuk and Tom Smith for assistance with statistical analyses, and Febrimarsa for introducing *Perkinsus* transformation to our studies. This work was supported by grants from the Gordon and Betty Moore Foundation (GBMF 4977, 4977.01) and King Abdullah University of Science and Technology (KAUST) (BAS/1/1020-01-01).

## MATERIALS AND METHODS

### Cell culture

*Perkinsus marinus* strain (ATCC 50983) was grown at 25 °C in ATCC medium 1886. Cells were typically subcultured by transfer of 200 μl of stationary-phase culture in 10 ml of medium in a T-25 culture flask every week for maintenance, or every three days if cells need to be used for experiments in exponential growth phase. *P. marinus* cultures were cryopreserved in ATCC medium 1886 media containing 10% DMSO, placed in a freezing container at −80°C for 24 h, and then moved to liquid nitrogen for long-term storage.

### Electroporation of *Perkinsus* cells

Plasmid DNA for transformation was prepared by mixing 2.5 μg linearised and 2.5 μg circular plasmid with 37 μl of 3R buffer (200 mM Na_2_HOP_4_, 70 mM NaH_2_PO_4_, 15 mM KCl, 150 mM HEPES pH: 7.3) and water to a final volume of 60 μl. *P. marinus* cells were grown to a cell density of OD600 = 0.4-0.5, corresponding to approximately 5-7×10^7^ cells and collected by centrifugation at 1000 x g, for 5 minutes at room temperature. The cell pellet was resuspended in the DNA-3R transformation mixture, and 10 μl of 1.5 mM CaCl2 was added to the suspension prior transfer into a 2 mm gap electroporation cuvette and electroporation by an Amaxa Nucleofector II electroporator, program D-023. Immediately after electroporation 0.5 ml of culture medium was added to the cuvette and all contents transferred to 6-well culture plates with a further 2.5 ml of growth media. Cells were allowed to recover at 25 °C for 48-72 hours prior additional experiments or start of drug selection.

### Golden Gate cloning

The cloning and domestication (removal of internal BsaI and BpiI sites) of genetic parts were generally performed as described in (Engler et al., 2014). The list of all PCR primers and plasmids constructed can be found in Supplementary Table S1. Restriction-ligation reactions were performed in 0.2 mL tubes containing the following: 200 ng of acceptor vector, 400 ng of each insert DNA component (be it a PCR product or plasmid), 5 U of the required restriction enzyme (BsaI/Eco31l, #ER0292, Thermo Fisher, or BpiI, ER1011, Thermo Fisher), and 400 U T4 DNA ligase (M0202, NEB), 1.5 μl T4 ligase buffer, 1.5 μl Bovine Serum Albumin (10 mg/ml) and water to a final volume of 15 μl. The reactions were performed in a thermocycler in one of two different reactions, first typically used for assembling more complex parts mixes. A longer reaction of 26 cycles of 3 minutes 37°C and 4 minutes 16°C followed by 5 minutes 50°C and 5 minutes 80°C. An alternative shorter cycling program consisted of 3 cycles of 10 minutes 37°C and 10 minutes 16°C followed by final incubation at 37°C for 10 minutes and 20 minutes at 65°C. For the shorter cycling program, the ligation buffer and BSA were replaced by 2 μl of Buffer G and 2 μl 10 mM ATP in a final volume of 20 μl, all enzyme concentrations were the same. The reaction mixture was added to DH5α competent cells (C2987H, NEB) and heat shock transformation performed according to the manufacturer’s recommendation. Transformed cells were selected on LB plates containing the appropriate antibiotic (Spectinomycin for L0, Kanamycin for L1 and Ampicillin for L2), and IPTG and X-gal for blue-white screening. Inserts were confirmed by Sanger sequencing. The sequences of all plasmids assembled and used in this study are available in Supplemental Table S1.

To test if the ‘PRDX’ promoter could drive expression of two genes, *luc* and *pac*, an overlap extension PCR was used to fuse parts. First, we amplified the reverse complement of luciferase and *moe* terminator sequences. Thereafter the plasmid ‘pRFW_L1_PAC3’ was used as a template to amplify the PRDX promoter fused to PAC and the *moe* terminator. These two PCR products were fused by a final PCR extension and cloned into the Level 1 position 1 acceptor vector for transfection, selection, and luciferase activity assay. The 2A sequences coupled to the *pac* gene and *moe* terminator sequences were synthesized (GeneArt, Thermo Fisher). The stop codon of mCherry was removed and overhangs were adjusted to be compatible to the 2A L0 part by PCR.

### Flow cytometry analysis

For FACS profiling, *P. marinus* cultures in stationary phase were diluted 1:2 with growth media or used undiluted if in exponential phase. Cells were filtered into 5 ml polystyrene tubes with a cell strainer cap (12×75mm) and 35 μm nylon mesh (Corning Falcon 352235) prior to analysis to remove cell aggregations. Cells were then analysed using an Aria III cell sorter with either a red laser (633 nm, 18mW) or a yellow-green laser (561 nm, 50 mW). For single-cell sorting, a 30 ml culture exponential growth was sorted at one cell per well in a 96-well plate containing conditioned ATCC medium 1886. The 96-well plates were incubated at 25°C to allow recovery and inspected weekly to monitor cell growth. After 4-8 weeks, cells were transferred to 6-well culture plates for further expansion of the clonal cell lines. Non-transformed *P. marinus* cells were always used as a negative control to set gating for single, non-fluorescence live cells. Data analysis was performed using FlowJo.

### Drug preparation, sensitivity assay, and selection

Stock solutions of puromycin (P8833, Merck) and blasticidin S-HCl (R21001, Thermo Fisher) were prepared by dissolving the drugs in sterile water at 50 mg/ml and 10 mg/ml, respectively. The solutions were then filtered with an 0.22-μm pore size MF-Millipore Membrane Filter and aliquoted to be stored at - 20°C. Bleomycin/zeocin (R25001, Invitrogen) were purchased as stock solution 100 mg/ml and was stored at - 20°C. *P. marinus* growth was assayed by OD600 using a Helios g Spectrophotometer with assays commenced at OD=0.15. Drugs were added to their test concentrations and aliquots taken and measured every second day over 15 days. All experiments were performed in triplicates.

For drug selection, post transformed cells were allowed to recover 3-4 days in 6-well plates before transferring them to T25 flasks in 10 ml of growth medium and selection drug (20 μg/ml puromycin, 200 μg/ml blasticidin S, or μg/ml bleomycin). Cultures resuspended in fresh media and selective drug each week following gentle pelleting (1000 x g, 5 min). Upon evidence of active cell growth (after approximately 25 days) culture passaging commenced as for wildtype cells.

### Luciferase activity assay

The luciferase ORF was amplified from Promega plasmid pGL4.11[luc2P] for use in Golden Gate assembly. For luciferase activity assays we typically used 5 ml of cell culture. We performed the luciferase activity assays according to the manufacturers’ protocol (#E2620, Bright-Glo assay kit, Promega) on cell cultures approximately 30 days post-transfection, or as soon as the cultures reached active growth. Cells were maintained with constant puromycin selection applied over the duration of measurement recordings. Non-transformed cells were used as negative control. Luminescence was recorded using GloMax plate reader (Promega). The analysis was performed on biological triplicates, except in four cases when only two cell lines were recovered after drug selection. For statistical analyses, the RLU values were normalized by culture density, log^10^ transformed and medians calculated of three technical replicates for each biological replicate of each promoter or control. The activity of each promoter was compared by one-way ANOVA analysis of variance followed by Dunnett’s multiple comparison test. Differences were statistically significant when *P* < 0.05. Prism 7 (GraphPad) was used for the statistical test.

### Genomic sequencing gDNA extraction, NGS library preparation, and sequencing

Clonal cell lines were selected by drug treatment and FACS sorting as described above. Genomic DNA was extracted using a DNeasy Blood and Tissue Kit (Qiagen), following the manufacturer’s instructions. gDNA quality was analysed by agarose gel and quantified using a Nanodrop. The gDNA was sheared with a Covaris E-Series ultrasonicator using the following program: Target BP (Peak) 550-600bp, Peak Incident Power (W) 105, Duty Factor 5%, Cycles per Burst 200, Treatment Time (s) 70. Multiplexed NGS libraries were prepared using the TruSeq PCR Free kit according to the manufacturer’s instructions (20015963, Illumina). The obtained libraries were sequenced using a Miseq 600 cycle kit to generate 150 bp and 300 bp paired-end reads to a depth of approximately 5-20X coverage. Raw data was de-multiplexed and quality filtered. Following this the genome of each clone was assembled using MaSuRCA 3.3.0 (Zimin et al., 2013) at standard settings. All transfectants were fluorescent: this allowed identification of scaffolds containing integrated plasmid sequence by BLAST using GFP or mCherry sequences as bait. After read mapping using bowtie2, reads mapped to these scaffolds were extracted using samtools and remapped for visual confirmation of plasmid integration (for the integration site sequence and bam files of the alignments see: Figshare DOI: 10.6084/m9.figshare.14611578). Substantial reorganization through indels of the transfected sequences was observed following alignment of genomic scaffolds and raw, untransformed plasmid sequences using MUSCLE 3.8.31 (Edgar, 2004).

### Western blot

Protein lysates were prepared from cell cultures approximately 8 weeks post transfection. The absorbance of the cell cultures was measured at 600 nm and then diluted to an optical density of 0.05 OD units per 10 μl. Samples for SDS-PAGE analysis was heated to 70°C for 10 minutes in 1x NuPAGE LDS sample buffer (NP0007, Thermo Fisher Scientific) with 2-Mercaptoethanol to a final concentration of 2.5%. Proteins were resolved by SDS-PAGE using NuPAGE 4-12% Bis-Tris Protein gels (Thermo Fisher Scientific) and electrotransferred to 0.2 μm pore size nitrocellulose membranes (Amersham Protran Supported, GE Healthcare). The membranes were blocked for 1 hour at room temperature using 5% (w/v) non-fat dry milk in either PBS or TBS containing 0.05% (w/v) Tween20. Membranes were thereafter probed with primary antibodies at 4°C for 16 hours with the following dilutions: 1:2000 anti-mCherry (ab167453, Abcam), 1:5000 anti-BiP, 1:1000 anti-Histone H3 (ab1791, Abcam) and 1:200 anti-Puromycin N-acetyltransferase 1HCLC (711421, Thermo Fisher Scientific). The secondary antibody, rabbit anti-goat peroxidase (115-035-003, Jackson Immunoresearch) was diluted 1:10000 for all blots except to detect Puromycin N-acetyltransferase, where the dilution was 1:5000. After a two-hour incubation at room temperature, the protein bands were visualised via chemiluminescence detection using SuperSignal West Pico substrate (Thermo Scientific). Images were recorded on a ChemiDoc MP+ imaging system (BioRad).

## Supplemental Information

**Figure S1.**
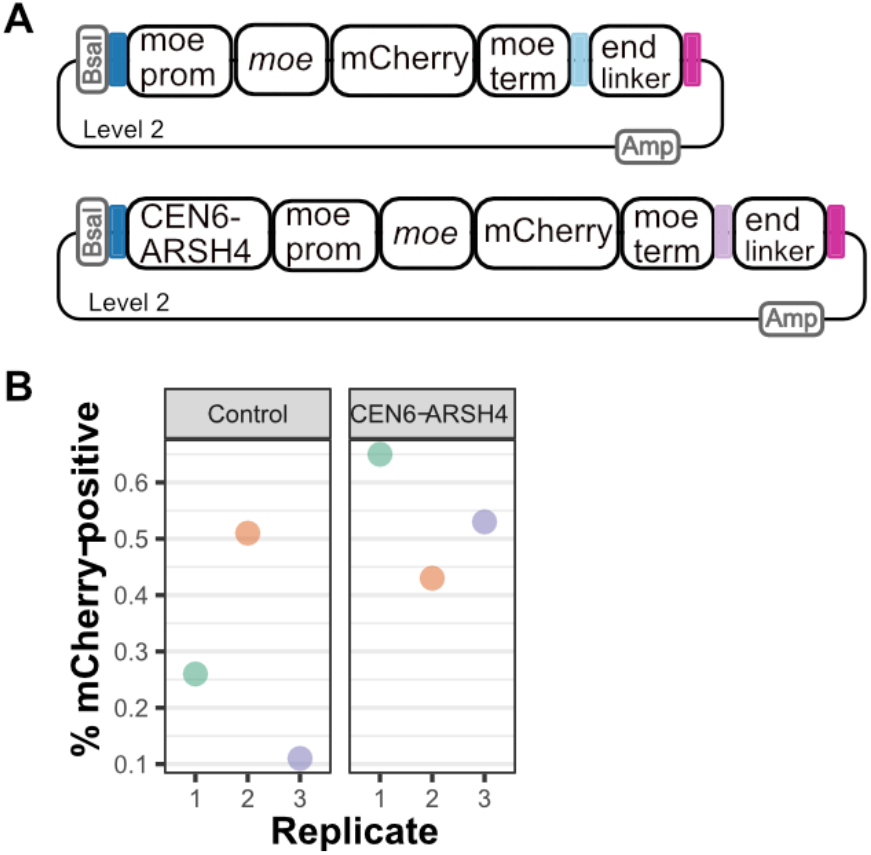
Test of plasmid retention in *P. marinus* using the CEN6-ARSH4 sequence. (**A**) Maps of MOE-mCherry expression plasmids either without or with the CEN6-ARSH4 sequence derived from a yeast artificial chromosome. (**B**) Percentage fluorescent *P. marinus* cells after mCherry-based sorting followed by14 days of growth without selection. No difference in plasmid retention was seen between the control (minus CEN6-ARSH4) and CEN6-ARSH4-containing plasmids by Student’s t-test of three replicates.

**Supplementary Table S1:** Primer and plasmid sequences.

